# Benchmarking bacterial genome-wide association study (GWAS) methods using simulated genomes and phenotypes

**DOI:** 10.1101/795492

**Authors:** Morteza M. Saber, Jesse Shapiro

**Affiliations:** Département de Sciences Biologiques, Université de Montréal, Montreal, QC, Canada

## Abstract

Genome Wide Association Studies (GWASs) have the potential to reveal the genetics of microbial phenotypes such as antibiotic resistance and virulence. Capitalizing on the growing wealth of bacterial sequence data, microbial GWAS methods aim to identify causal genetic variants while ignoring spurious associations. Bacteria reproduce clonally, leading to strong population structure and genome-wide linkage, making it challenging to separate true “hits” (i.e. mutations that cause a phenotype) from non-causal linked mutations. GWAS methods attempt to correct for population structure in different ways, but their performance has not yet been systematically evaluated. Here we developed a bacterial GWAS simulator (BacGWASim) to generate bacterial genomes with varying rates of mutation, recombination, and other evolutionary parameters, along with a subset of causal mutations underlying a phenotype of interest. We assessed the performance (recall and precision) of three widely-used univariate GWAS approaches (cluster-based, dimensionality-reduction, and linear mixed models, implemented in PLINK, pySEER, and GEMMA) and one relatively new whole-genome elastic net model implemented in pySEER, across a range of simulated sample sizes, recombination rates, and causal mutation effect sizes. As expected, all methods performed better with larger sample sizes and effect sizes. The performance of clustering and dimensionality reduction approaches to correct for population structure were considerably variable according to the choice of parameters. Notably, the elastic net whole-genome model was consistently amongst the highest-performing methods and had the highest power in detecting causal variants with both low and high effect sizes. Most methods reached good performance (Recall > 0.75) to identify causal mutations of strong effect size (log Odds Ratio >= 2) with a sample size of 2000 genomes. However, only elastic nets reached reasonable performance (Recall = 0.35) for detecting markers with weaker effects (log OR ∼1) in smaller samples. Elastic nets also showed superior precision and recall in controlling for genome-wide linkage, relative to univariate models. However, all methods performed relatively poorly on highly clonal (low-recombining) genomes, suggesting room for improvement in method development. These findings show the potential for whole-genome models to improve bacterial GWAS performance. BacGWASim code and simulated data are publicly available to enable further comparisons and benchmarking of new methods.

**Author summary:** Microbial populations contain measurable phenotypic differences with important clinical and environmental consequences, such as antibiotic resistance, virulence, host preference and transmissibility. A major challenge is to discover the genes and mutations in bacterial genomes that control these phenotypes. Bacterial Genome-Wide Association Studies (GWASs) are family of methods to statistically associate phenotypes with genotypes, such as point mutations and other variants across the genome. However, compared to sexual organisms such as humans, bacteria reproduce clonally meaning that causal mutations tend to be strongly linked to other mutations on the same chromosome. This genome-wide linkage makes it challenging to statistically separate causal mutations from non-causal false-positive associations. Several GWAS methods are currently available, but it is not clear which is the most powerful and accurate for bacteria. To systematically evaluate these methods, we developed BacGWASim, a computational pipeline to simulate the evolution of bacterial genomes and phenotypes. Using simulated genomes, we found that GWAS methods varied widely in their performance. In general, causal mutations of strong effect (*e.g.* those under strong selection for antibiotic resistance) could be easily identified with relatively small samples sizes of around 1000 genomes, but more complex phenotypes controlled by mutations of weaker effect required 3000 genomes or more. We found that a recently-developed GWAS method called elastic net was particularly good at identifying causal mutations in highly clonal populations, with strong linkage between mutations – but there is still room for improvement. The BacGWASim computer code is publicly available to enable further comparisons and benchmarking of new methods.

## Introduction

Recent progress in sequencing technologies and consequently rapid expansion of bacterial genomic data repositories have provided enormous opportunities to identify the genomic elements underlying clinically, environmentally and industrially important bacterial phenotypes and their evolutionary responses to changing environmental circumstances. Such discoveries could immensely improve our knowledge of the molecular mechanisms of important microbial phenotypes such as antibiotic resistance and virulence, thus contributing to the development of new drugs, vaccines and antibiotics.

By identifying statistical associations between genotype and phenotype, Genome Wide Association Studies (GWASs) can be used to dissect the genetic components of any measurable and heritable phenotype in an unbiased hypothesis-free manner. In humans, GWASs have been used to investigate genotype-phenotype association since the early 2000s, leading to the discovery of more than 149,000 trait-associated genomic markers in thousands of publications (1). In bacteria, the GWAS approach can be traced back to a 2005 attempt to unravel the genomic elements responsible for transforming harmless *Neisseria meningitidis* into a lethal pathogen causing cerebrospinal meningitis using Multi Locus Sequence Typing (MLST) (2). The difficulties and limitations of bacterial GWASs imposed by population structure were appreciated soon thereafter (3). Nevertheless, over the past decade GWASs applied to Single Nucleotide Polymorphisms (SNPs) and *K-*mers (i.e. DNA words of length *K*) in microbial genomes has identified mutations and genes associated with antibiotic resistance (4–10), cancer (11), virulence (2,12,13) and host preference (14). In contrast to human GWAS, however, bacterial association mapping is technically challenging due to the unique characteristics of bacterial populations, and optimal approaches for bacterial GWASs have yet to be established.

The objective of a GWAS method is to maximize statistical precision and power, in order to identify true causal genomic elements while ignoring spurious associations. To do so, GWAS methods must overcome confounding factors which are particularly acute in bacterial populations. The two main confounding elements in bacteria are genome-wide linkage disequilibrium interrupted by homologous recombination tracts, and strong population structure resulting from clonal expansions.

Genome-wide linkage disequilibrium (LD) leads to type I errors (false positives) in GWAS tests because linked non-causal mutations may hitchhike on the same genomic background (“clonal frame”) as a causal mutation. A naïve GWAS approach will find the entire set of linked mutations to be associated with the phenotype. In bacterial species such as *M. tuberculosis* that are virtually non-recombinogenic, most of the genome is in complete LD, posing a major risk of type I error. In humans and other sexual species, LD is broken down by homologous recombination every generation, allowing GWAS tests to map the causal variants to a small genomic region. In bacteria, LD may span the entire genome, complicating fine mapping of causal variants. Even in bacterial species with relatively high rates of homologous recombination (*e.g. S. pneumoniae*), LD may still extend across the entire chromosome.

Population structure refers to a situation in which subpopulations have systematic differences in allele and phenotype frequencies (15,16). This can result in spurious associations between genotypes and phenotypes due to shared ancestry rather than causal associations. To control for the confounding effect of population structure, microbial GWAS tools have adapted approaches already used in human GWASs, including cluster-based techniques (6,12,17), dimensionality reduction (18–22), Linear Mixed Models (LMMs) (4,8,10,23,24), and recently, artificial intelligence using whole-genome models (25–27). Although each of these approaches has been successful to varying extents, population structure is still a challenge in microbial GWASs and no gold standard solution has been established.

The power of a GWAS to identify causal variants underlying a phenotype is influenced by several other factors, including the sample size and the distribution of effect sizes. In human GWASs, most of the detected causal variants underlying complex phenotypes have odds ratio (OR) < 1.5 (28) due to the polygenicity of the traits and the fact that many human phenotypes of interest are disease traits that have been largely shaped by neutral evolution rather than strong selection. In bacteria, however, many phenotypes of interest such as antibiotic-resistance or host-association tend to be shaped by recent positive selection. Therefore, the genomic elements controlling bacterial traits are expected to have larger effect sizes (17). For example, mutations conferring antibiotic resistance in *Mycobacterium tuberculosis* (29) and *Steptococcous pneumoniae* (6) tend to have large effect sizes (OR > 10). However, it has yet to be investigated whether smaller effect sizes are detectable, and if so, by which methods.

To determine best practices for microbial GWAS, it is essential to compare current GWAS methods in terms of their performance across a range of realistic effect sizes, recombination rates, and sample sizes. For this purpose, here we developed a simulation platform called BacGWASim which simulates bacterial genomes along a defined phylogenetic tree to capture mutation and recombination events in a clonal population structure. BacGWASim is tunable for a variety of evolutionary parameters in order to simulate a wide range of bacterial species. It then simulates bacterial phenotypes based on adjustable values of heritability, number of causal variants, and their effect sizes. The major different classes of GWAS methods currently in use were then evaluated in terms of their precision and power to identify true causal variants in the simulated bacterial populations. In summary, we provide an extensible framework for simulating the evolution of bacterial genotypes and phenotypes, and for benchmarking new GWAS approaches as they become available.

## Results

### Simulating bacterial genomes and phenotypes

To systematically benchmark bacterial GWAS approaches, we first developed an appropriate simulator of bacterial genomes and phenotypes, BacGWASim (Methods; Figure 1a). The simulator starts from a real reference genome with gene annotations and then allows this genome to evolve along a user-defined or simulated phylogenetic tree capturing the population structure (Figure 1b). Phenotypes are simulated according to a heritability function of causal SNPs, with user-defined effect sizes. Realistic sources of noise, such as sequencing error and read mapping, are also included. Although other evolutionary parameters (*e.g.* mutation rates) are tunable in the simulation, here we varied three key parameters with a likely effect on GWAS performance: sample size, recombination rate, and effect size of causal mutations. We began with the genome sequence and phylogenetic tree of a well-studied species, *Streptococcus pneumoniae* (although the user could alternatively choose to simulate a phylogeny *de novo* using a birth-death process). A range of genome wide linkage disequilibrium (LD) was then simulated to approximate the range of LD observed in bacterial species with high, moderate and low recombination rates. While not an exact match, these simulations approximate the LD landscapes of *S. pneumoniae, E. coli*, and *M. tuberculosis*, respectively (Figure 2). We will begin by focusing on the high-recombining, low LD simulations to explore the effects of sample and effect sizes on GWAS performance, and then revisit the effects of LD to conclude. In all simulations, phenotype heritability was set to one, with an equal number of cases and controls in each dataset. The BacGWASim codes are publicly available at: https://github.com/Morteza-M-Saber/BacGWASim.

**Figure 1.**
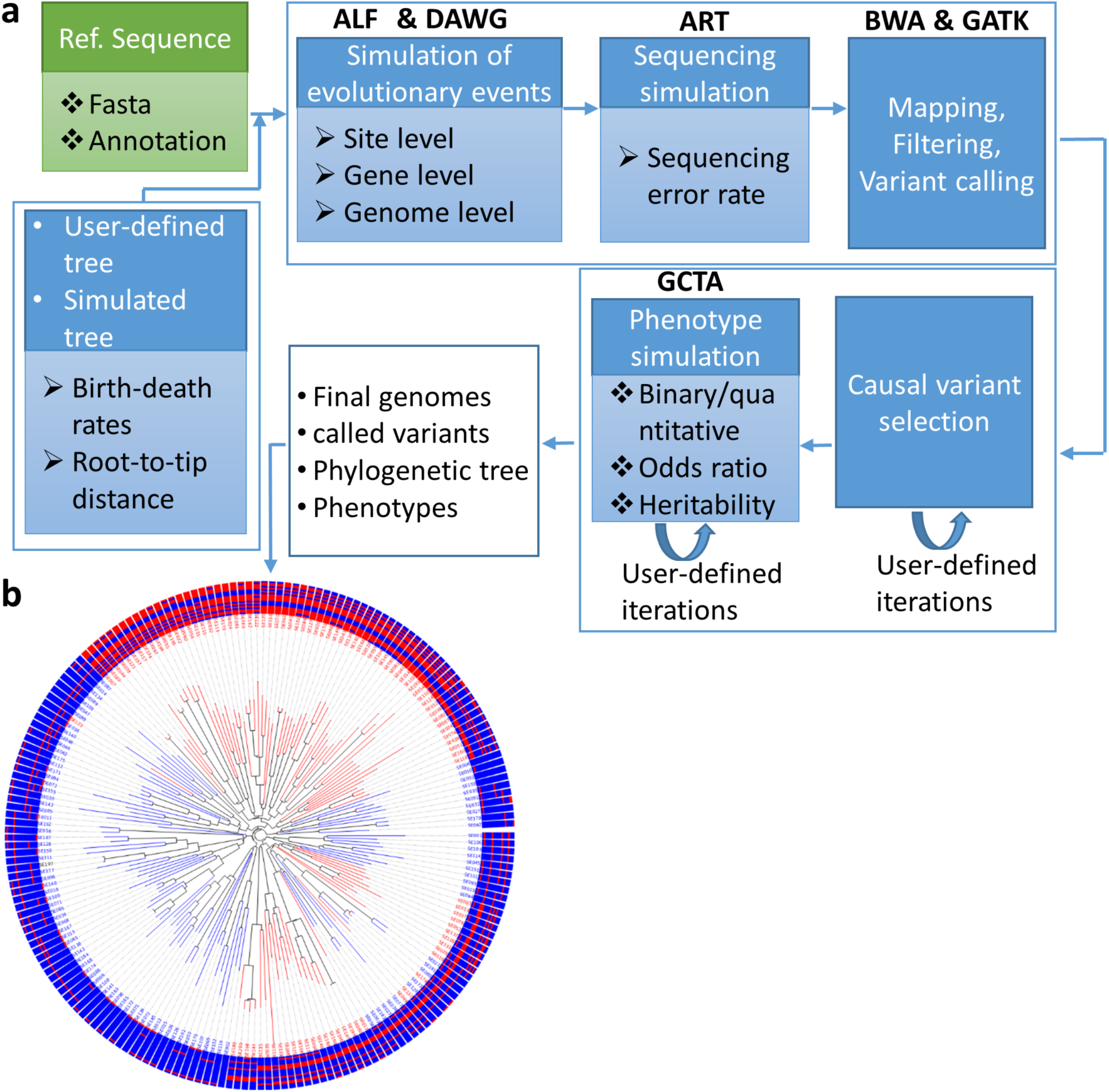
Overview of the bacterial GWAS simulation (BacGWASim) pipeline. (a) Taking a bacterial genome and annotations as input, evolutionary events are simulated across the branches of the phylogenetic tree (either user-defined or simulated based on a birth-death model), producing the final genomes and simulated markers as output. A phenotype is then assigned to each simulated sample based on the presence/absence of a set of causal SNPs. (b) Binary phenotypic states, shown as red or blue branches, mostly cluster in monophyletic subpopulations of closely-related individuals due to population structure. The external rings denote the allele (red or blue) at each of the simulated causal loci (shown as concentric rings).

**Figure 2.**
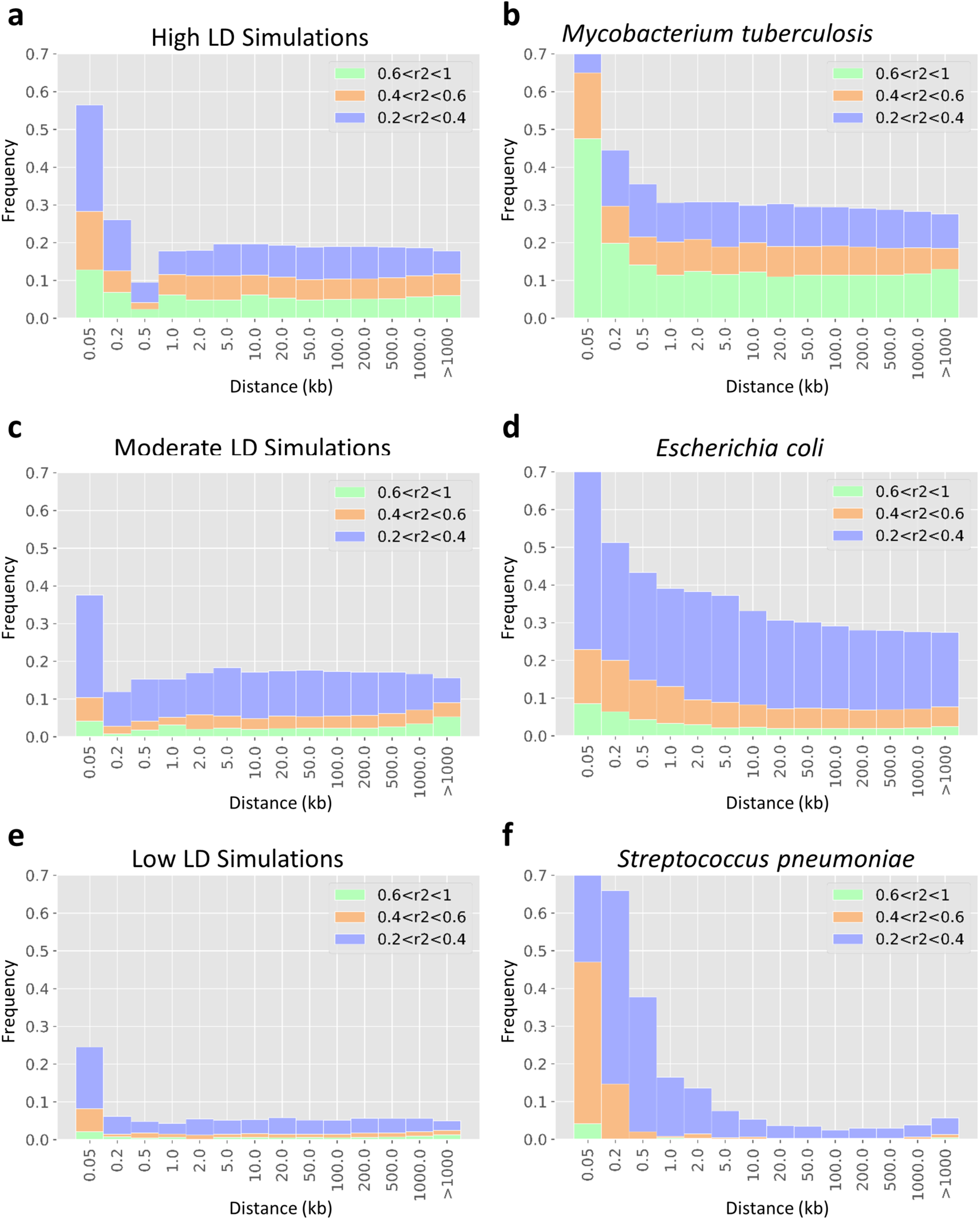
Landscape of genome-wide linkage disequilibrium (LD) in bacterial species compared to BacGWASim simulations. The distribution of linkage, measured as r^2^ scores binned into three categories, is shown as a function of distance in the genome, in units of kilobase pairs, within genomes of simulated (left panels) and real (right panels) bacteria.

### GWAS power to detect variants of large effect reaches a plateau at 3000 genomes

Early GWASs were able to identify causal variants of large effect with relatively small sample sizes (on the order of 100 genomes) likely because the phenotypes investigated were under strong positive selection (4,9,12,14,17). However recent advances in high-throughput sequencing technologies now makes it feasible to sequence thousands of genomes. To evaluate the effect of increasing sample size on the power to detect causal mutations within a range of effect sizes, we measured the performance of bacterial GWAS methods on a range of sample sizes. Simulating a high-recombining, low LD population (Figure 2e), the elastic net whole-genome model implemented in pyseer was consistently amongst the most accurate across the range of sample sizes, with F1 score ranging between 0.44 to 0.60 (Figure 3a) and fewer than 50 false positives out of 100,000 tested markers (Figure 3c). The linear mixed model (LMM) implemented in GEMMA and clustering approach implemented in plink also showed low rates of false positives comparable to elastic nets (Figure 3c); however, they had lower recall (Figure 3b). FaST-LMM implemented in pyseer, despite its high recall (Figure 3b) achieved lower F1 scores due to its relatively high false positive rates (Figure 3c). With relatively small sample sizes (<1000), all methods except for elastic net showed poor performance in detecting causal variants with low effect sizes.

**Figure 3.**
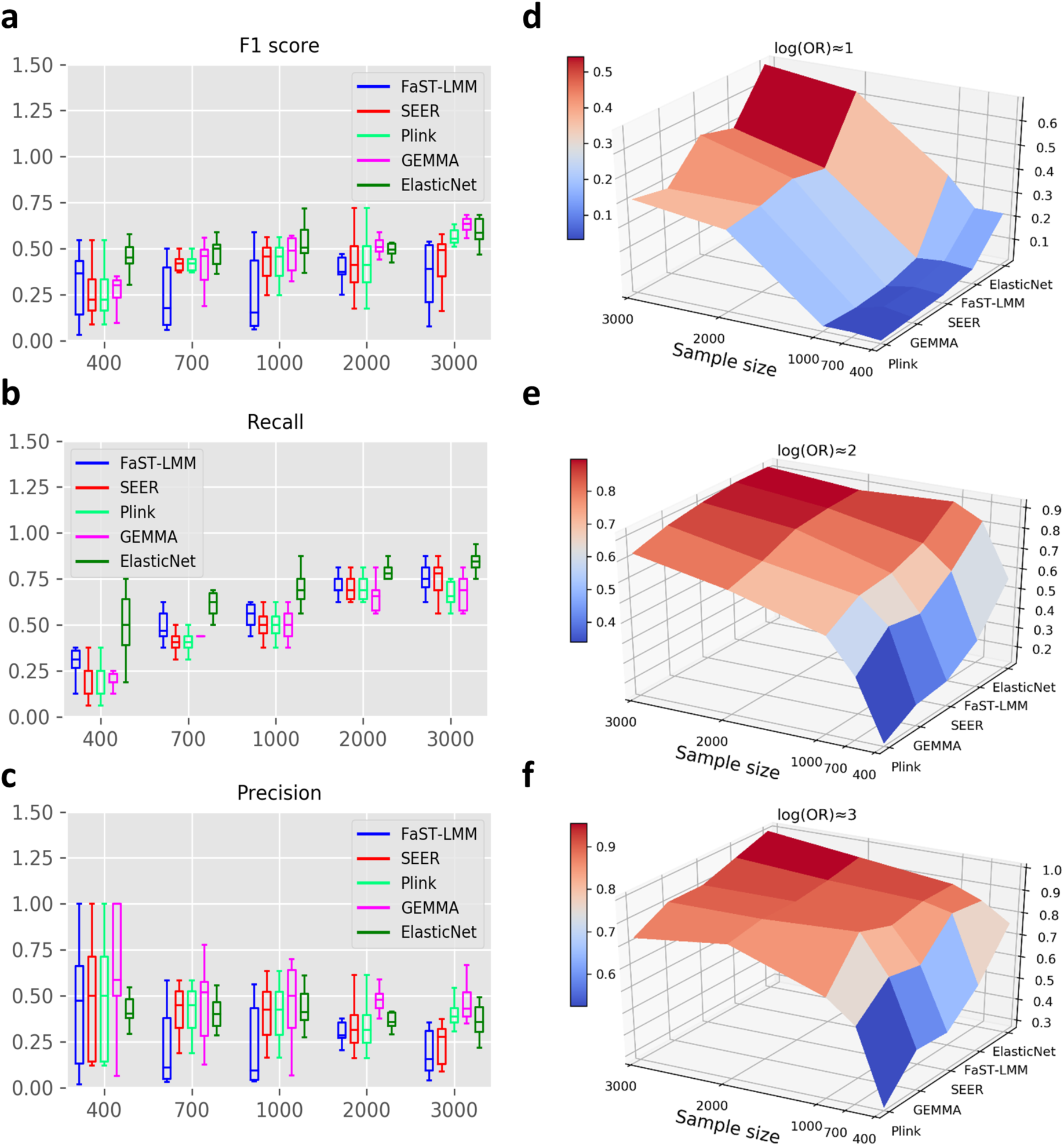
Performance of GWAS methods in simulations with low LD. The boxplots on the left (a, b, c) show the median and interquartile range, averaged across a range of effect sizes (log (OR=1 to 3) The surface plots on the right (d, e, f) show recall rates in separate categories of effect sizes. (a) F1 score of bacterial GWAS methods across a range of sample sizes under low LD. (b) Recall rates across the range of sample sizes. (c) Precision rates across the range of sample sizes.

A general pattern observed across all methods is that after a rapid surge in recall rate by increasing the sample size to 1000, the power improvement slows down and reaches a plateau around 3000 samples (Figure 3b). Breaking down these comparisons by effect size shows that, as expected, causal markers with low effect size (log (OR) ≈1) are the most sensitive to sample size. The power to detect such variants is < 0.1 in univariate methods and < 0.35 in elastic nets in samples sizes below 1,000. Power improves to 0.37-0.48 for univariate methods and 0.68 for elastic nets at sample size of 3,000 (Figure 3d). In contrast, recall rate (power) for causal markers with higher effect sizes (log (OR) ≈2 and log (OR) ≈3) is more uniform across methods and reaches a maximum of 0.78 in univariate models and in elastic nets, with only 1,000 genomes sampled. Beyond 1,000 samples, the increase in recall tends to be small, reaching a plateau around 3,000 samples for most methods (Figure 3e and 3f).

### Correcting for population structure

We next investigated how GWAS power varies across the four methods to correct for population structure: univariate models including cluster-based approaches, dimensionality reduction, and linear mixed models, and the whole-genome model using elastic nets. In univariate models, the correlation between each of the markers (SNPs or K-mers) and the desired phenotype is investigated separately. Therefore, the covariance between the markers due to population structure needs to be included explicitly. Whole-genome models, on the other hand, include all the genetic variants at once and by this means, covariance between the variants are implicitly included in the analysis. Our findings show that the whole-genome elastic net model has superior power relative to univariate models in controlling for population structure and genome-wide LD, especially in small sample sizes (Figure 3d, e, f) and when there is strong linkage across the genome, as discussed below. First, we consider each of the univariate methods separately.

#### Cluster-based approaches

One of the classic methods for controlling the confounding effect of population structure is to identify clusters of related individuals within the overall population and then test for association conditional on these subpopulations. Subpopulations can be inferred using a variety of methods (30–33) and then a weighted association test is performed for each genomic marker across the defined clusters (e.g. with the Cochran–Mantel–Haenszel test). The proportion of population structure captured in this approach, however, depends on the threshold used for clustering. Choosing a strict threshold for clustering will improve the precision of the test but at the expense of reducing the recall score. To measure the effect of the choice of clustering threshold on the power of cluster-based methods, we performed linkage agglomerative clustering based on IBS distances implemented in PLINK across a range of thresholds. The threshold was defined as the maximum number of individuals allowed to group in one cluster. By relaxing the threshold, a larger number of samples with lower genetic similarity are included in each cluster and therefore, population structure will not be fully captured. In contrast, setting a very strict threshold will make the association test conservative by over-correcting for population structure.

Using simulated datasets of 3,000 genomes with low LD (Figure 2e), we varied the population structure correction from strict (maximum number of two individuals per cluster) to weak (up to a hundred individuals per cluster). As expected, going from strict to weak correction improves recall (from 0.56 to 0.75) at the expense of precision, which drops from 0.57 to 0.38 (Figure 4a). The choice of clustering threshold significantly affects the ability to detect variants of low effect (log OR≈1), but has little effect on variants of larger effect (Figure 4b). In the absence of established standards to choose a clustering threshold, our results suggest that the choice may not matter for the detection of large-effect variants, but could significantly bias the detection of markers with low effect sizes.

**Figure 4.**
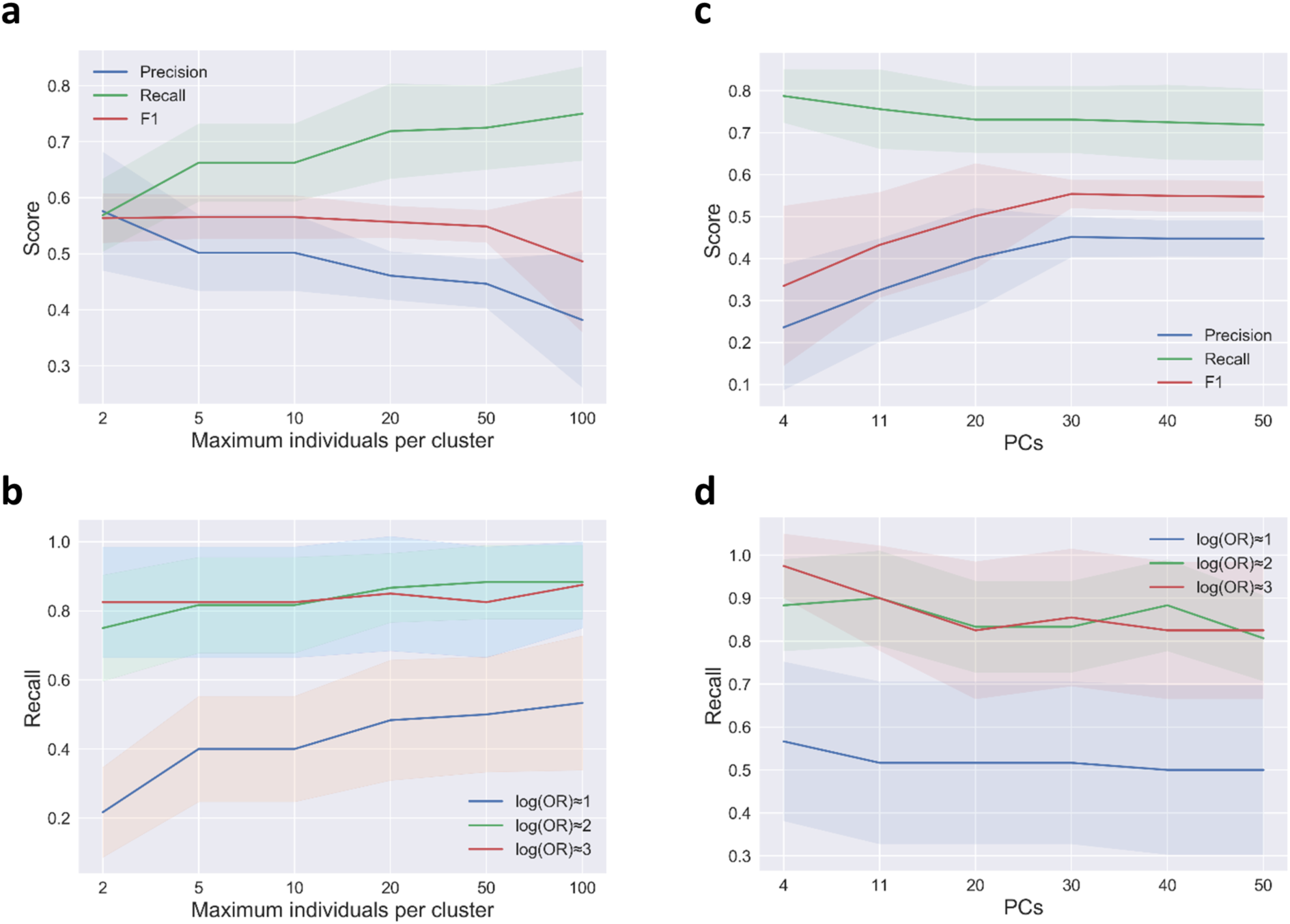
GWAS performance varies according to different levels of correction for population structure in a dataset of 3,000 genomes simulated under low LD. (a) Effect of the clustering threshold in terms of the maximum number of individuals per cluster on GWAS. (b) GWAS power (recall) across a range of clustering thresholds for identifying causal markers within separate categories of effect sizes. (c) Effect of variation in the number of included principal components as covariates used for correction. (d) GWAS power (recall) across a range of included principal components for detecting causal markers within separate categories of effect sizes.

#### Dimensionality reduction

In dimensionality reduction, a relatedness matrix of samples is projected onto a lower number of components, and then a certain number of these components are chosen as fixed effect factors in the linear regression (or logistic regression in the case of binary phenotypes) to account for population structure. The two most popular methods for dimensionality reduction in GWAS are principal component analysis (PCA) of variance-standardized relationship matrix based on SNP data (33) and metric multidimensional scaling (MDS) of genetic similarities based on pairwise distance matrix constructed using the shared k-mer content between all samples (18). The former has been widely used in eukaryotic GWAS studies where there is low variation in pan-genome size, while the latter is specifically designed for bacterial GWAS and uses an alignment-free k-mer based approach to estimate genetic similarities based on core and accessory genome distances. Like cluster-based approaches, the proportion of population structure controlled in this approach depends on the number of included components as covariates in the regression analysis. Although there are some recommended methods for determining the optimal number of included components such as visual estimation using the scree plot (34), the number of included components is subjective and is a matter of sensitivity-specificity tradeoff. Including more components is likely to improve the precision of GWAS and reduce Type I error caused by population structure, at the expense of decreasing the recall score.

To evaluate the effect of the number of included components in dimensionality reduction-based bacterial GWAS, we tested the SEER method implemented in pyseer on a simulated high-recombining dataset of 3,000 samples (Figure 2e). According to the scree plot, 4 or 11 are good choices of the number of components to be included in the regression analysis, as there are major and minor drops after these number of dimensions (Supplementary Figure 1). However, the F1 score reaches its maximum after inclusion of 30 dimensions (Figure 4c). By increasing the number of components from 4 to 30, the precision significantly improved from 0.23 to 0.45, then reaches a plateau. Meanwhile, the recall score is relatively unaffected (dropping from 0.79 to 0.73), leading to total improvement of F1 score (from 0.33 to 0.55) with an increasing number of components. As expected, recall is better for markers of high or medium effect, but recall does not vary much with increasing components for any category of effect size (Figure 4d). In general, our results indicate that the choice of included components significantly affects the performance of dimensionality reduction-based stratification correction, and the lack of the standard protocol to identify the optimum number of included principal components can limit the application of this method.

#### Linear mixed models (LMMs)

The LMM is an extension of linear regression, which allows the inclusion of both fixed and random effects as covariates. By using a kinship matrix to model the variance of a random effect, LMMs consider the genetic relationships between all samples rather than selecting a proportion of the population structure (as in the cluster-based and dimensionality-reduction approaches described above) and has been shown to control type I error without loss of power (35). GEMMA (36) and FaST-LMM (37) are two popular LMM-based GWAS methods and have been used in recent bacterial GWASs, either as standalone methods, or as implemented in BugWAS (23), DBGWAS (24), or pyseer (34).

To compare the power of LMM-based GWAS methods, we tested the performance of GEMMA and FaST-LMM implemented in pyseer on the same simulated high-recombining dataset of 3,000 samples (Figure 2e). Pyseer provides the option to construct the kinship matrix using either variant-based genetic distances or by extracting patristic distances from the phylogeny, while GEMMA recommends using the centered genotype matrix. Averaged across effect sizes, GEMMA is the most efficient method to control for type I errors caused by population structure and results in the highest F1 score (Figure 5a). In FaST-LMM, the phylogeny-based correction for population structure outperforms the genotype matrix, mainly due to a boost in precision (Figure 5a). However, in this case the ‘true’ phylogeny (known from the simulation) was used, but must be estimated in real applications. Therefore, the accuracy of a phylogenetic correction might be lower depending on the choice of methods to construct the phylogenetic tree. FaST-LMM has slightly higher recall (power) than GEMMA, and its advantage was most pronounced for variants with low effect sizes (Figure 5b). All methods had high and nearly equal power (> 0.85) in identifying causal variants with high effect sizes (log (OR) ≈2 and log (OR) ≈3). However, for detecting low effect-size causal variants (log (OR) ≈1), FaST-LMM has higher power (Figure 5b).

**Figure 5.**
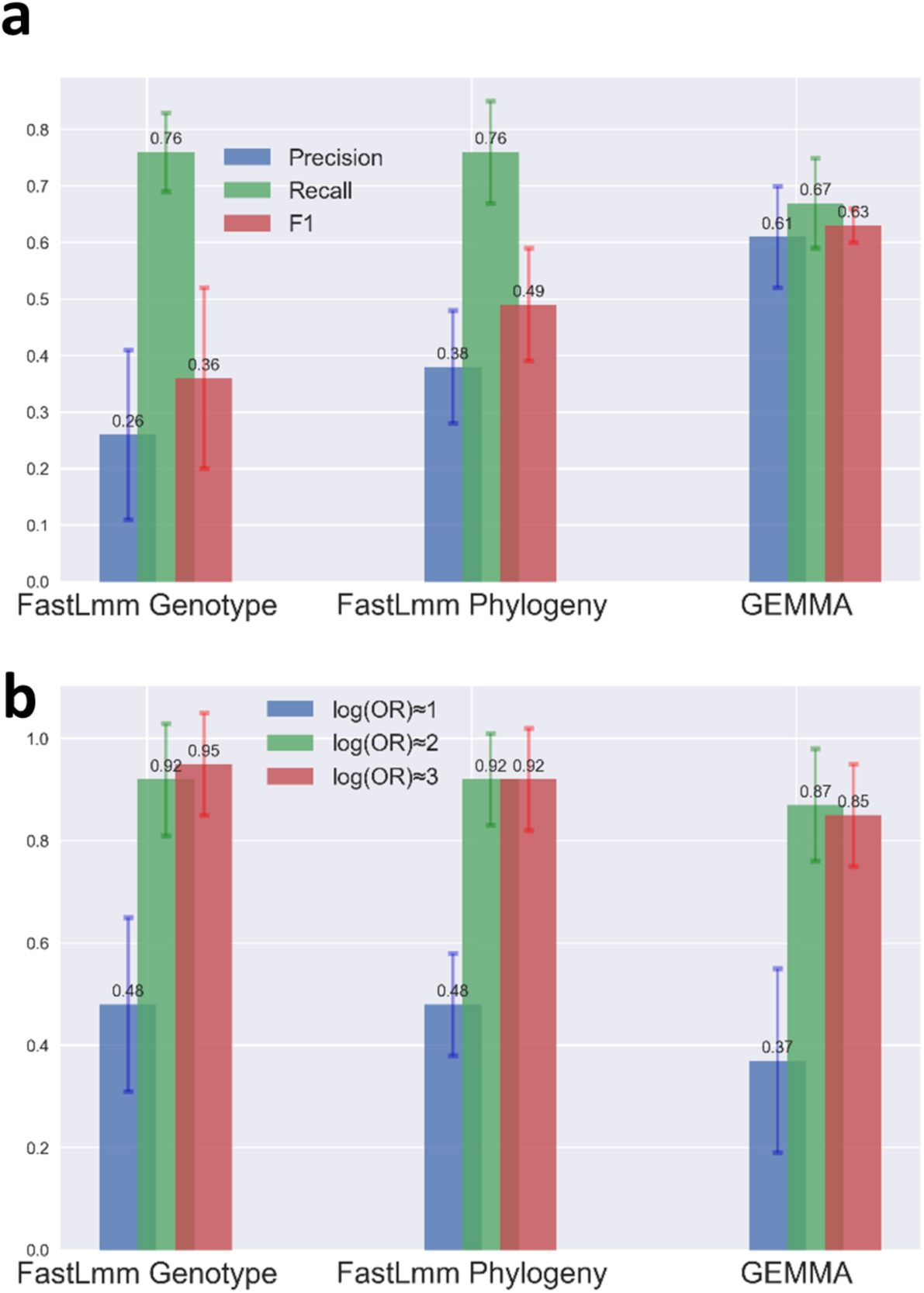
Performance of LMM-based GWAS methods in a dataset of 3,000 simulated genomes under low LD. (a) Comparison of different implementations of linear mixed models in terms of precision, recall and F1 scores. (b) Comparison of the recall (power) of different implementations of linear mixed models in identifying causal markers within separate categories of effect sizes.

### Current GWAS methods perform poorly under moderate to high LD

Higher genome-wide LD is expected to reduce the precision of association testing, because hitchhiking non-causal mutations are identified as false positives. To assess the influence of LD on GWAS performance, we tested each method across a range of simulated datasets with low, moderate, or high genome-wide LD (Figure 2). In general, the elastic net implemented in pyseer considerably outperforms other methods at moderate or high LD (Figure 6). At low LD, elastic nets perform similarly to GEMMA, the best univariate method (Figure 6a). At high LD, elastic nets achieve ∼75% power, compared to ∼60% in univariate methods (Figure 6b). All methods suffer a loss of precision with increasing LD, but elastic nets retain the highest precision at high LD (Figure 6c). Still, the median precision of elastic nets at high LD is ∼8%, suggesting significant room for improvement. The FaST-LMM approach using the genotype-matrix for population structure correction was most severely affected by LD, with precision dropping from 0.36 at low LD to <0.001 at high LD (Figure 6c). On the other hand, FaST-LMM with the phylogeny-based correction showed comparable results to the best performing univariate-model methods. Amongst the univariate models, the mixed model approach implemented in GEMMA tended to perform somewhat better than others at moderate or high LD (Figure 6a). Q-Q plots further indicate GEMMA to be the best performing method across the range of LD among the univariate models (Supplementary Figure 2).

**Figure 6.**
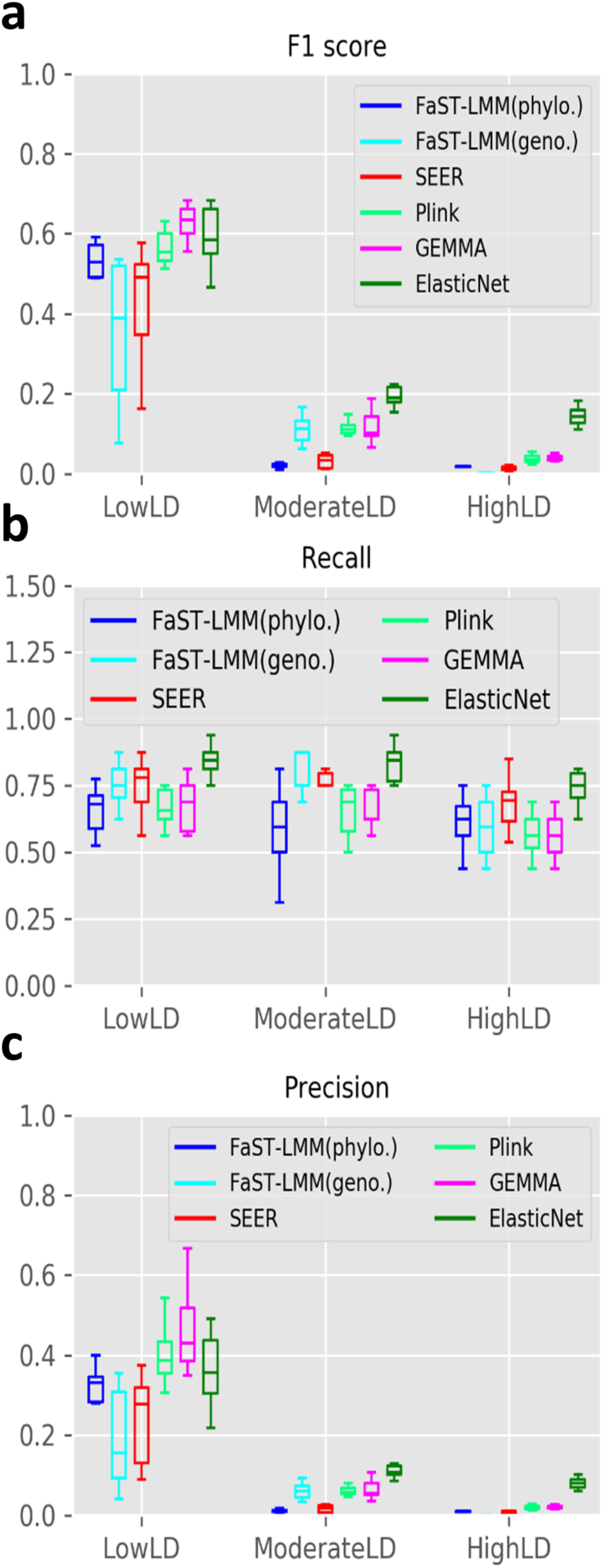
Effect of LD on GWAS performance. (a) F1 score (b) recall and (c) precision of GWAS methods in datasets of 3000 genomes with low, moderate and high genome-wide LD levels.

## Discussion

We developed a platform to simulate bacterial genomes and phenotypes based on the emergence and evolution of causal variants along a phylogenetic tree. The simulator is tunable for relevant evolutionary parameters. Although it was designed for benchmarking GWAS tools, it could also be used for other applications in bacterial population genomics and epidemiology. Here we focused on how GWAS method performance is affected by sample size, recombination (linkage disequilibrium), and causal variant effect sizes. We only considered point mutations in the core genome as causal variants, but this could be extended to causal variants in the pangenome (*i.e.* gene presence/absence), which can be identified with k-mer approaches like SEER. Although a full exploration of core and pangenome associations is beyond the scope of the current work, we expect our major findings to hold when causal variants in the pangenome are included. Our evolutionary model is also ‘neutral’ in that it does not consider positive selection on causal variants (*e.g.* antibiotic resistance mutations that have a selective advantage). Future versions of BacGWASim could explicitly model an increased replication rate (clonal expansion) of bacteria encoding causal variants, perhaps more realistically capturing the evolution of certain phenotypes.

Our results confirmed the suspicion that currently popular GWAS methods perform poorly when applied to bacteria with relatively clonal population structures yielding moderate or high LD (*e.g. E. coli* or *M. tuberculosis*). In such clonal populations, fine-mapping of causal mutations may be impossible and identification of phenotype-associated lineages may be the best possible outcome (23). More encouragingly, causal variants can be detected with relatively high power and precision in higher-recombining populations, akin to *S. pneumoniae*. These results highlight the importance of assessing the LD landscape of the target organisms before deciding on a sample size and GWAS design. It also suggests significant room for improvement in GWAS method development, particularly for highly clonal bacteria. Promising methods include phyC (9) and treeWAS (15) which detect homoplastic (convergent) mutations along a clonal phylogeny, providing strong control over population structure given accurate phylogenetic tree. Due to the high computational burden of these tests, especially with large sample sizes (35), we did not evaluate these methods here. Moreover, although homoplasies do occur in our simulations (Figure 1b), their rate is not explicitly controlled and thus their impact is hard to assess. In the future, BacGWASim could be extended to explicitly model homoplastic mutations and to assess the performance of homoplasy-based methods.

Of the GWAS methods evaluated here, the elastic net whole-genome model generally performed best, followed by the mixed model approach implemented in GEMMA. The clustering approach implemented in PLINK also performed well, but varied significantly depending on the clustering threshold, which can be challenging to optimize. Elastic nets implemented in pyseer also provide the possibility to perform a GWAS on k-mers or unitigs, whereas this is not as easily done in GEMMA or PLINK. Although elastic nets had the highest precision of the methods tested, there is significant room for improvement, as mentioned above.

Our results also help explain the success of early bacterial GWASs with low sample sizes. We found that, in a high-recombining population, a sample size ∼1,000 is sufficient to reliably detect causal variants of strong effect (log OR >= 2) with high power (>0.80). Such strong effect sizes may be common for antibiotic resistance mutations, and other variants under strong positive selection. However, samples sizes >3,000 will likely be needed to detect variants of lower effect (log OR ∼1), which may be more common for more ‘complex’ phenotypes with lower heritability. Of course, the sample size required to achieve a desired power will vary depending on the recombination rate and population structure of the organism of interest. BacGWASim thus provides a tool for study-specific power calculations.

In an age of a rapidly growing array of options for performing GWASs, we hope that our results are instructive in quantifying general trends and sources of bias, and that our simulation platform can continue to be used to benchmark novel methods as they appear. For example, machine learning has recently been used in bacterial GWAS, with successful application in highly clonal species such as *Mycobacterium tuberculosis* (25–27,38). These methods have the advantages of being highly versatile with high computational efficiency. Although not all artificial intelligence methods are currently packaged for the purpose of bacterial GWAS, our findings suggest great potential for whole-genome models to further fine-tune bacterial GWAS. Our simulation platform also provides a means to evaluate and benchmark their performance against the traditional methods as they emerge.

## Methods

### Overview of the Bacterial GWAS Simulator (BacGWASim)

BacGWASim was developed with the goal of simulating a range of evolutionary parameters which can potentially confound microbial genome-wide association studies. In this first release, we primarily focused on simulating population structure and genome-wide linkage disequilibrium, and evaluating GWAS to identify SNPs underlying a phenotype. BacGWASim starts with a bacterial whole genome and its annotations and simulates a population across a pre-defined or simulated phylogenetic tree (Figure 1a). A binary or continuous phenotype is then assigned to each genome in the population based on the emergence and evolution of randomly chosen set of causal variants with a user-defined range of effect sizes.

BacGWASim simulate bacterial genotypes and phenotypes in three main steps: (1) generating a phylogenetic tree, (2) evolving genomes along the phylogenetic tree, and (3) simulating phenotypes.

### Generating the phylogenetic tree

A neutral model of speciation-extinction developed by Genhard (39) implemented in Artificial Life Framework (ALF) (40) was used for the simulation of phylogenetic trees. This model allows for simulation of bacterial populations across a wide range of birth rates (λ), death rates (µ) and branch length distributions. A custom phylogenetic tree can also be provided by the user to provide the option for simulation of scenarios inferred from real bacterial populations.

### Simulation of genome evolution

For simulation of realistic genomic content, BacGWASim accepts any bacterial whole genome and the corresponding feature annotations as starting input. The elements of the genome are then extracted and classified in two groups of protein coding genes and intergenic regions (defined as sequence not annotated as a CDS). These two categories are often under distinct evolutionary constraints and their evolution is best captured by different evolutionary parameters. Artificial Life Framework (ALF) was used for the simulation of protein coding gene and DAWG v.2 for the intergenic regions (41). These implementations allowed for simulation of three categories of genomic events: 1) site-level events including codon and nucleotide substitution rates, insertion and deletion rates, and rate variation across sites, 2) gene-level events including gene deletion, duplication and gene fission/fusion, and 3) genome-level events including inversion, translocation and recombination through horizontal gene transfer. The resulting sequences of coding and intergenic regions at the tips of phylogenetic tree are then combined while accounting for gene loss and transfer. Synthetic sequencing reads with Illumina-specific sequencing errors were generated for each simulated sample using ART next-generation sequencing read simulator (42). These synthetic reads were then mapped to the reference genome using BWA (43) with default setting and the simulated variants were called using GATK Haplotype caller by setting ‘ploidy’ to 1 (44).

### Phenotype simulation

After genome simulation, a binary or continuous phenotype was then simulated for each member of population based on an additive genetic model implemented in GCTA (45). For this purpose, a user-defined number of causal variants, minor allele frequency cutoff, and range of effect sizes in units of odds ratio are used to randomly select a set of “true” causal variants from the pool of simulated markers. The phenotype labels are then simulated according to the presence/absence of the causal variants in each genome, user-defined values of heritability, and prevalence of the desired phenotype.

### Benchmark datasets and methods of Bacterial GWAS

Bacterial genomes were simulated within ranges of three features of interest, keeping other parameters constant:

- **Sample size**: bacterial populations with a range of 400 to 3000 sampled genomes were simulated to evaluate the effect of sample size on GWAS.
- **Recombination rate**: Recombination rates in highly-recombining *Streptococcus pneumoniae* estimated by Chewapreecha et al. (46) (average ρ/θ ≈0.20) were used for simulation of highly recombining populations. Moderate- and low-recombining populations were respectively simulated with ρ/θ ratios of 0.1 and 0.001.
- **Effect size distribution**: Eighteen causal markers with odds ratio (effect sizes) of 2, 3, 4, 7, 10, 11, 15 and 20 (Natural logarithm in the range of 1 to 3) with minor allele frequency >0.1 were randomly chosen for phenotype simulation. The linkage disequilibrium between the selected markers were measured using bcftools (47) and markers with r^2^ > 0.6 were discarded.

To measure the range of LD in real bacterial species, genome data were retrieved from the CARD database (48), SNPs were called using Snippy (49) and linkage levels were measured in 1,000 markers using Haploview (50) (Figure 2).

In every set of simulations, 100,000 randomly selected markers with minor allele frequency >0.01 were retained for GWAS analysis. An equal number of markers was selected in each simulation to make them comparable without any confounding effect of multiple testing across replicates. For each genome simulation, ten sets of randomly chosen markers were then used to generate ten replicate phenotype simulations. Using the called variants and phenotype labels as benchmark datasets, we then used the following methods to perform GWAS:

- **Plink v1.9** using linkage agglomerative clustering based on pairwise identity-by-state (IBS) distances for population structure correction, and using Bonferroni corrections for multiple tests (33).
- **SEER** implement in pyseer v 1.2.0 using multi-dimensional scaling of pairwise k-mer based genetic distances to correct for population structure, using unique patterns to estimate significance threshold, and by removing the markers tagged with “af-filter”, “bad-chisq”, “pre-filtering-failed”, “lrt-filtering-failed”, “perfectly-separable-data”, “firth-fail” and “matrix-inversionerror” after the analysis (18,34).
- **Fast-LMM implemented in pyseer 1.2.0 using pairwise variant-based distances** to correct for population structure, using unique patterns to estimate significance threshold, and by removing the tagged markers mentioned above (as for SEER) after the analysis (34,37).
- **Fast-LMM implemented in pyseer 1.2.0 using phylogeny-based patristic distances** to correct for population structure, using unique patterns to estimate significance threshold, and by removing the tagged markers mentioned above after the analysis (34,37).
- **GEMMA v.98** using pairwise variant-based genetic distances to correct for population structure and setting the option ‘gk’ to 1 for generating the relationship matrix.
- **Elastic-net whole-genome model implemented in pyseer 1.2.0**, by setting the alpha value to 1, without sequence reweighting, and using Bonferroni correction for multiple tests, based on the second round of model fitting and feature selection (34).

The performance of each GWAS method was assessed based on the mean values of precision, recall and F1 scores, and the corresponding standard deviations across ten replicate simulations for each parameter combination, where:

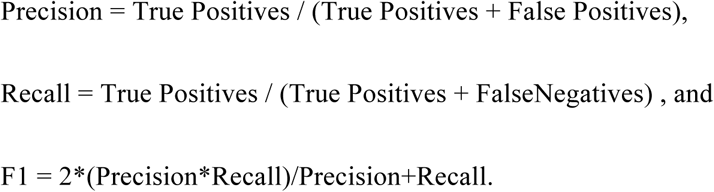

## Acknowledgements

This work was supported by a Genome Canada and Génome Québec grant to BJS. We are indebted to John Lees for useful discussions and valuable advice during the preparation of this manuscript.

## Supporting information

**Supplementary figures S1 and S2.** A document containing all the supplementary figures referenced in the text. (PDF)

## Author Contributions

Conceptualization: Morteza M. Saber, B. Jesse Shapiro.

Formal analysis: Morteza M. Saber.

Funding acquisition: Jesse Shapiro.

Investigation: Morteza M. Saber, B. Jesse Shapiro.

Methodology: Morteza M. Saber, B. Jesse Shapiro.

Project administration: Jesse Shapiro.

Software: Morteza M. Saber.

Visualization: Morteza M. Saber.

Writing– original draft: Morteza M. Saber, B. Jesse Shapiro.

Writing – review & editing: Morteza M. Saber, B. Jesse Shapiro.

