## Supplementary Figures for "Benchmarking bacterial genome-wide association study (GWAS) methods using simulated genomes and phenotypes"

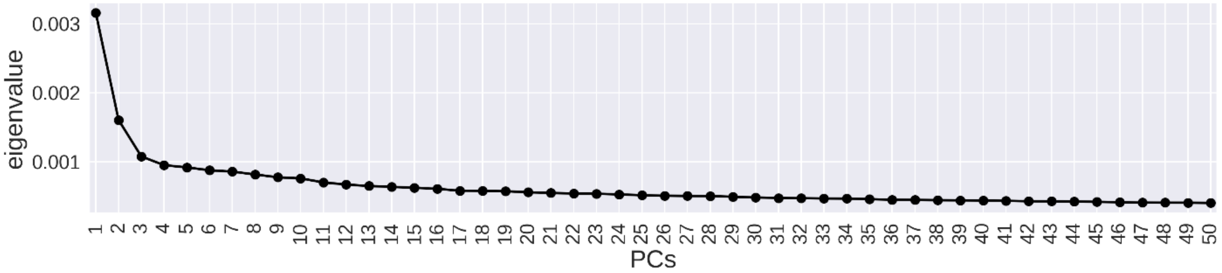


*Figure S1. Scree plot of multi-dimensional scaling analysis.*

There are major and minor drops after four and eleven dimensions indicating these values to be reasonable choices for the number of included components in the logistic regression analysis-based GWAS.

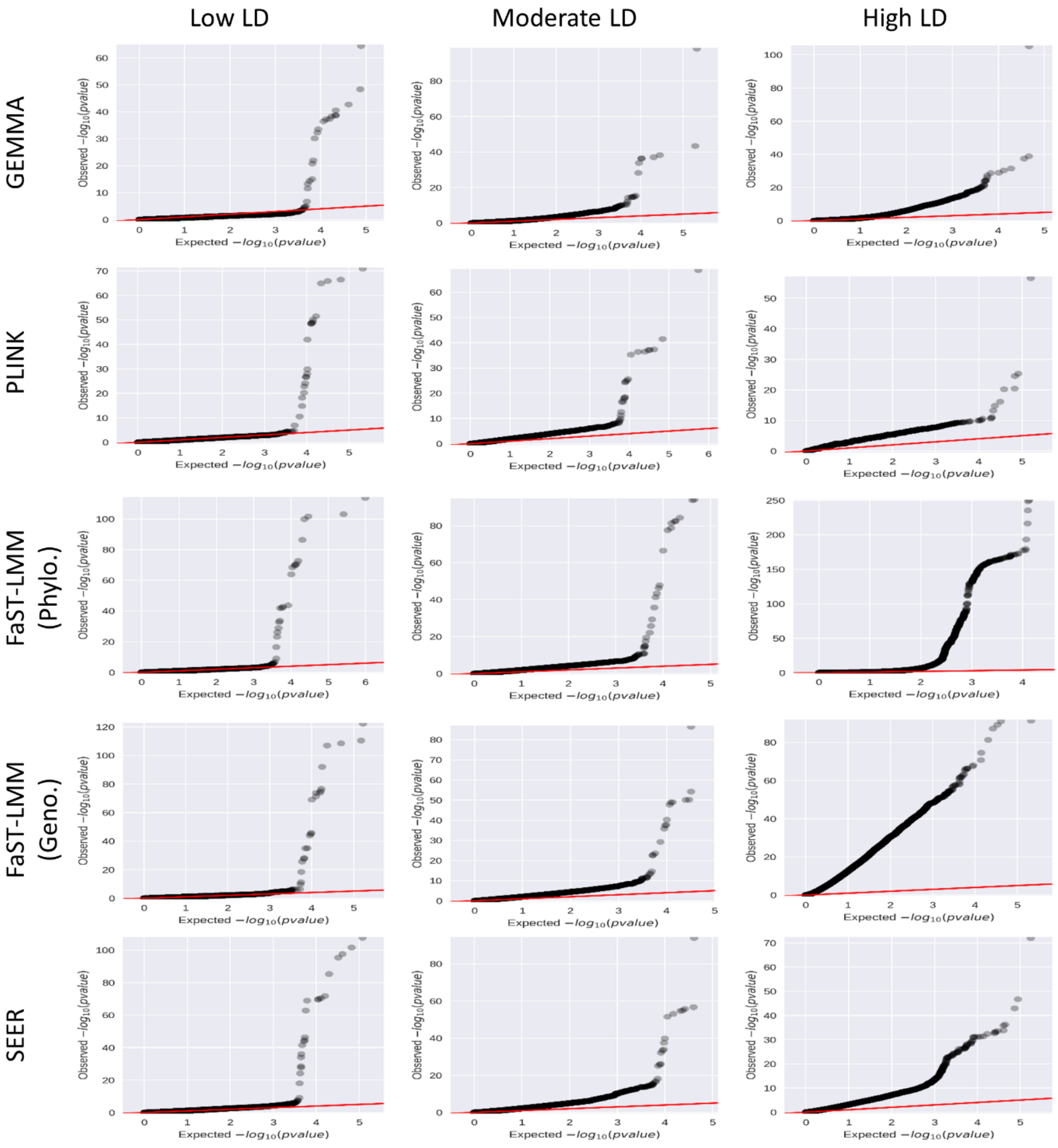


*Figure S2. Q-Q plots of GWAS analysis using univariate models.*

Q-Q plots of GWAS analysis in low, moderate and high LD datasets using GEMMA, the clustering approach in PLINK, FaST-LMM implemented in pyseer using phylogeny-based distances, FaST-LMM implemented in pyseer using variant-based distances, and dimensionality-reduction approach in SEER.
